# Aging alters the metabolic flux signature of the ER unfolded protein response *in vivo* in mice

**DOI:** 10.1101/2021.04.14.439896

**Authors:** Catherine P. Schneider, Lucy Peng, Samuel Yuen, John Halstead, Hector Palacios, Edna Nyangau, Hussein Mohammed, Naveed Ziari, Ashley E. Frakes, Andrew Dillin, Marc K. Hellerstein

## Abstract

Age is a risk factor for numerous diseases, including neurodegenerative diseases, cancers, and diabetes. Loss of protein homeostasis is a central hallmark of aging. Activation of the endoplasmic reticulum unfolded protein response (UPR^ER^) includes changes in protein translation and membrane lipid synthesis. Using stable isotope labeling, a “signature” of the UPR^ER^ *in vivo* in mouse liver was developed by inducing ER stress and measuring rates of both proteome-wide translation and *de novo* lipogenesis. Several changes in protein synthesis across ontologies were noted with age, including a more dramatic suppression of translation under ER stress in aged mice as compared to young mice. Binding immunoglobulin protein (BiP) synthesis rates and mRNA levels were increased more in aged than young mice. *De novo* lipogenesis rates decreased under ER stress conditions in aged mice, including both triglyceride and phospholipid fractions. In young mice, only a significant reduction was seen in the triglyceride fraction. These data indicate that aged mice have an exaggerated response to ER stress, which may indicate that the aging renders the UPR^ER^ less effective in resolving proteotoxic stress.

## Main

Loss of protein homeostasis, or proteostasis, is a central hallmark of aging and may explain why certain diseases become manifest as organisms grow older^1^. Proteostasis involves coordination of the synthesis of new proteins, quality control of the proteome and adaptive mechanisms to reduce unfolded and misfolded proteins and prevent abnormal protein aggregation^2^. Proteins are synthesized in the endoplasmic reticulum (ER) and chaperones to aid in the proper folding of newly synthesized proteins and to assist when protein misfolding occurs. Accumulation of misfolded proteins in the ER stimulates the unfolded protein response (UPR^ER^), an integrated set of adaptations that clear misfolded protein aggregates and either restore more normal proteostasis or ultimately eliminate affected cells through apoptosis^3,4^. The UPR^ER^ consists of three downstream pathways initiated by inositol-requiring enzyme-1 (IRE1), PKR-like ER kinase (PERK), and activating transcription factor 6 (ATF6), all of which are anchored in the ER membrane. ER-localized binding immunoglobulin protein (BiP), also identified as glucose-regulated protein 78-kD (GRP78), is one of the responders to misfolded proteins in the ER and acts as a regulator of the UPR^ER 5^. Downstream effects include global suppression of protein translation, with the exception of key proteins involved in a rescue response such as chaperones and lipogenic proteins^6^. If ER stress is unable to be resolved, cells undergo apoptosis^7^. Unmitigated ER stress may be a central component of many diseases, including metabolic diseases such as fatty liver disease and insulin resistance^8,9^.

In addition to aiding in restoration of proteostasis through halting global protein translation, the UPR^ER^ initiates ER membrane expansion through incorporation of fatty acids into the membrane to accommodate for aggregating proteins and chaperones recruited to assist in disaggregation or refolding^10^. Added ER surface may also help with the synthesis of necessary compensatory equipment, such as nascent proteins and lipids. The source of these lipids incorporated into hepatocyte ER was previously unknown, however, was recently discovered to be mobilized free fatty acids from adipose tissue during ER stress and not from local *de novo* lipogenesis^11^. Tunicamycin induced ER stress in mice leads reduction of lipogenic gene expression is reduced in livers of mice under ER stress^12,13^ and of *de novo* lipogenesis^11^. Alterations in metabolism and ER membrane expansion are crucial, yet poorly understood metabolically, as elements of the ER stress response^9,14–16^. Age induced metabolic changes and effect on ability to handle ER stress are especially poorly understood.

Although the UPR^ER^ has been implicated to decline with age in C. elegans^17^, among other organisms, it is not fully understood how age induced shifts in metabolism may impair an organisms’ ability to handle proteotoxic stress. Furthermore, proteins involved in the UPR may continue to be rapidly translated whereas translation of other proteins will be suppressed through phosphorylation of eIF2α by PERK^18,19^. Thus, measurement of protein fluxes provides a powerful tool for identifying UPR^ER^ regulators and signatures. In this study, we measured both proteome-wide replacement rates and *de novo* lipogenesis (DNL) through stable isotope labeling. We defined a flux “signature” of the unfolded protein response in mice liver, which revealed the proteins potentially involved in the rescue response of the UPR^ER^. Heavy water labeling in this experiment allowed measurement of newly synthesized proteins and fatty acids, such as palmitate, under induced ER stress, and the incorporation of newly synthesized palmitate into both phospholipids and triglycerides^21^. Phospholipids are especially of interest due to the thought that they may be incorporated into ER membranes under times of ER stress^22,23^. It is unknown how these ER stress induced metabolic fluxes change in aged mice, and can provide insight into how proteostasis breaks down with age.

## Results

In order to determine how induction of the unfolded protein response differs with age, both 12-week-old and 80-week-old male mice (n=5 per condition) were treated with 1 mg/kg tunicamycin once per day over a 4-day treatment period to induce a state of chronic ER stress. Tunicamycin inhibits N-linked glycosylation, leading to the accumulation of misfolded proteins^20^. ^2^H_2_O (deuterated or heavy water) administration began at the time of the initial tunicamycin treatment. Proteins synthesized after tunicamycin treatment incorporate deuterium-labeled amino acids, whereas pre-existing proteins will not have ^2^H label in covalent C-H bonds of amino acids, enabling the measurement of proteins that were newly synthesized during the period of exposure to tunicamycin. Mice were given deuterated water to also label nascent lipids *in vivo* (figure S1). Response to tunicamycin induced activation of the UPR^ER^ was characterized by proteome wide changes in translation and through *de novo* lipogenesis of palmitate incorporated into isolated phospholipid or triglyceride fractions.

### Proteome wide changes in translation signatures with initiation of the UPR

Through measuring deuterium incorporation into newly synthesized proteins by tandem mass spectrometric analysis, the fractional synthesis rates for proteins translating during the treatment period were measured. Key UPR^ER^ proteins, including protein disulfide isomerases, BiP, endoplasmin, and calreticulin, continued to be translated at higher rates than most global protein translation at 96 hours post initial tunicamycin treatment (figure 1a). However, global protein translation began to recover by 96 hours (figure 1b). To further understand the data, proteins were organized by their KEGG-pathways to calculate pathway specific rates of protein translation (figure 1c). By normalizing the synthesis rates in mice challenged with tunicamycin to control DMSO treated mice, the fold-change in protein synthesis rate by KEGG-pathway was determined. Under chronic ER stress conditions, protein processing in the ER was the most upregulated ontology at 2.6-fold higher under ER stress conditions as compared to control. *Fatty acid degradation* and *PPAR signaling* were two of the most suppressed ontologies. Other ontologies were mostly unaffected, with slight increase in overall translation.

**Figure 1:**
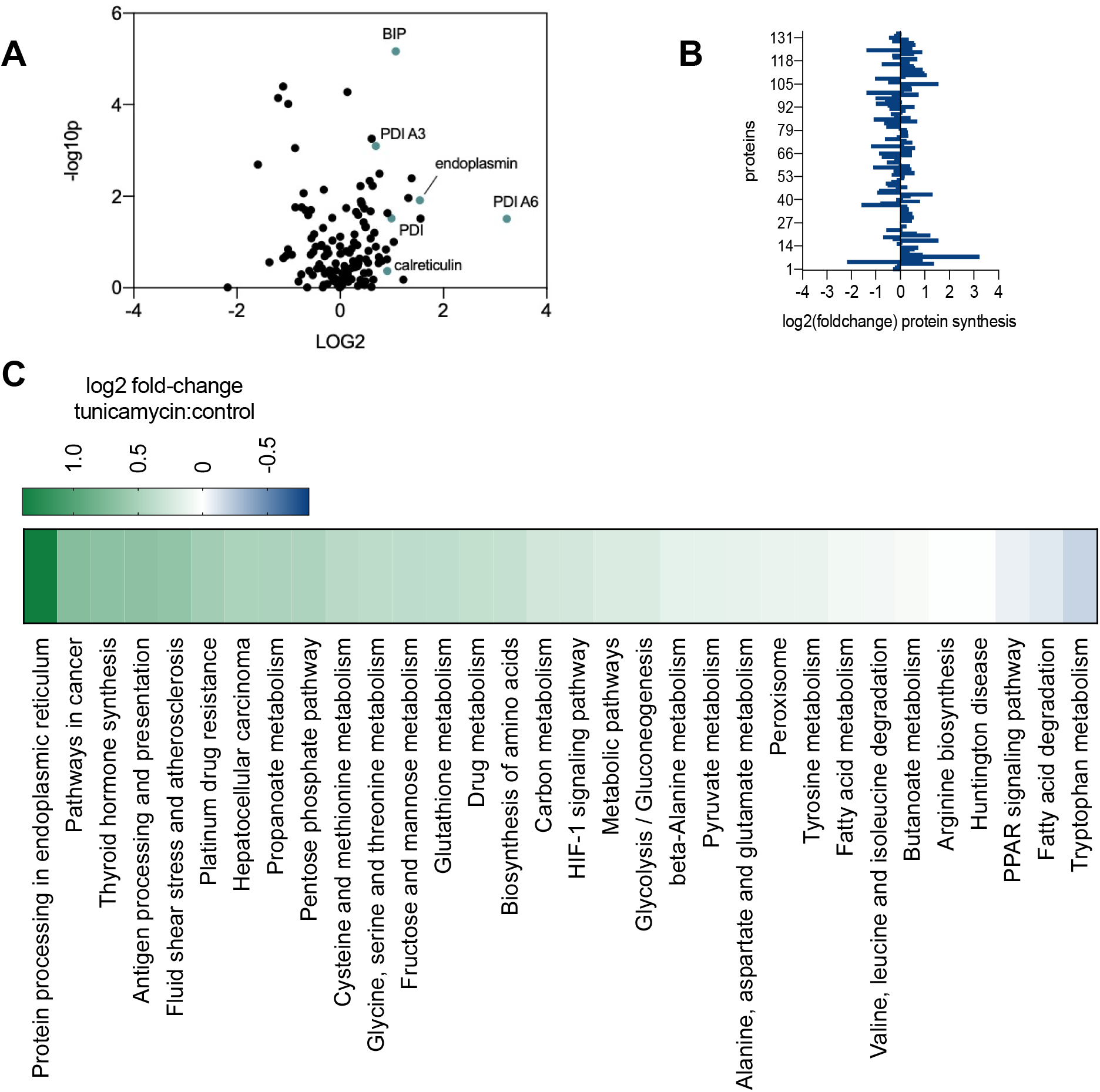
(a) Volcano plot of all hepatic proteins for which fractional synthesis rates were measured (n=136) in 12-week-old mice. Points expressed as log2 fold-change tunicamycin treated/control on x-axis and - log10(p-value), obtained from 2-tailed t-test, on y-axis. (b) log2 fold-change of individual protein translation rates of tunicamycin treated/control. (c) KEGG-pathway analysis for fractional synthesis rates of proteins from 12-week-old mice tunicamycin treated/control. n = at least 5 proteins per pathway.

### Age induced changes to characterized UPR signature

Aged mice exhibited a strikingly polarized response to chronic ER stress compared to their younger counterparts (figure 2a-b). Aged mice experienced broad inhibition of protein translation, with most ontologies being suppressed for protein synthesis. Proteins of the ontology *protein processing in the ER* remained as highly upregulated in the young mice, with a 2.6-fold increase compared to tunicamycin challenged young mice. Ontologies pertaining to lipid metabolism, including *PPAR signaling, fatty acid metabolism*, and *fatty acid degradation* were more suppressed in the aged mice as compared to young challenged with tunicamycin (figure 2b). Overall, when challenged with tunicamycin, aged mice showed much lower rates of translation across most ontologies (figure 3). In contrast, those ontologies that were upregulated remained as highly translated as the younger mice. When compared directly, under ER stress conditions most ontologies in aged mice were significantly suppressed in their synthesis of translation, with the exception of higher rates of synthesis of *protein processing in the ER*.

**Figure 2:**
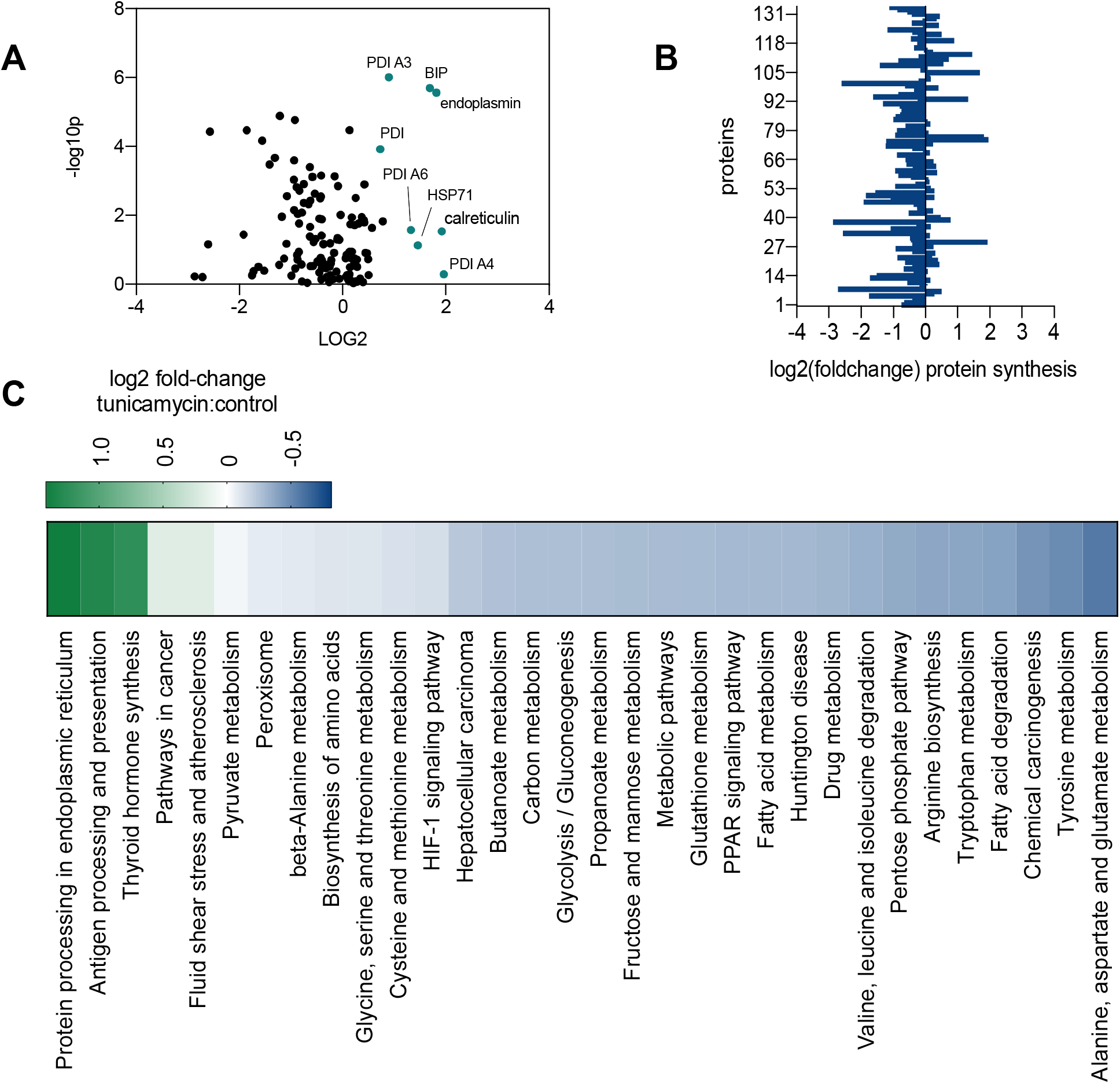
(a) Volcano plot of all proteins for which fractional synthesis rates were measured (n=136) in 80-week-old mice. Points expressed as log2 fold-change tunicamycin treated/control on x-axis and - log10(p-value), obtained from 2-tailed t-test, on y-axis. (b) log2 fold-change of individual protein translation rates of tunicamycin treated/control. (c) KEGG-pathway analysis for fractional synthesis rates of proteins from 80-week-old mice tunicamycin treated/control. n = at least 5 proteins per pathway.

**Figure 3:**
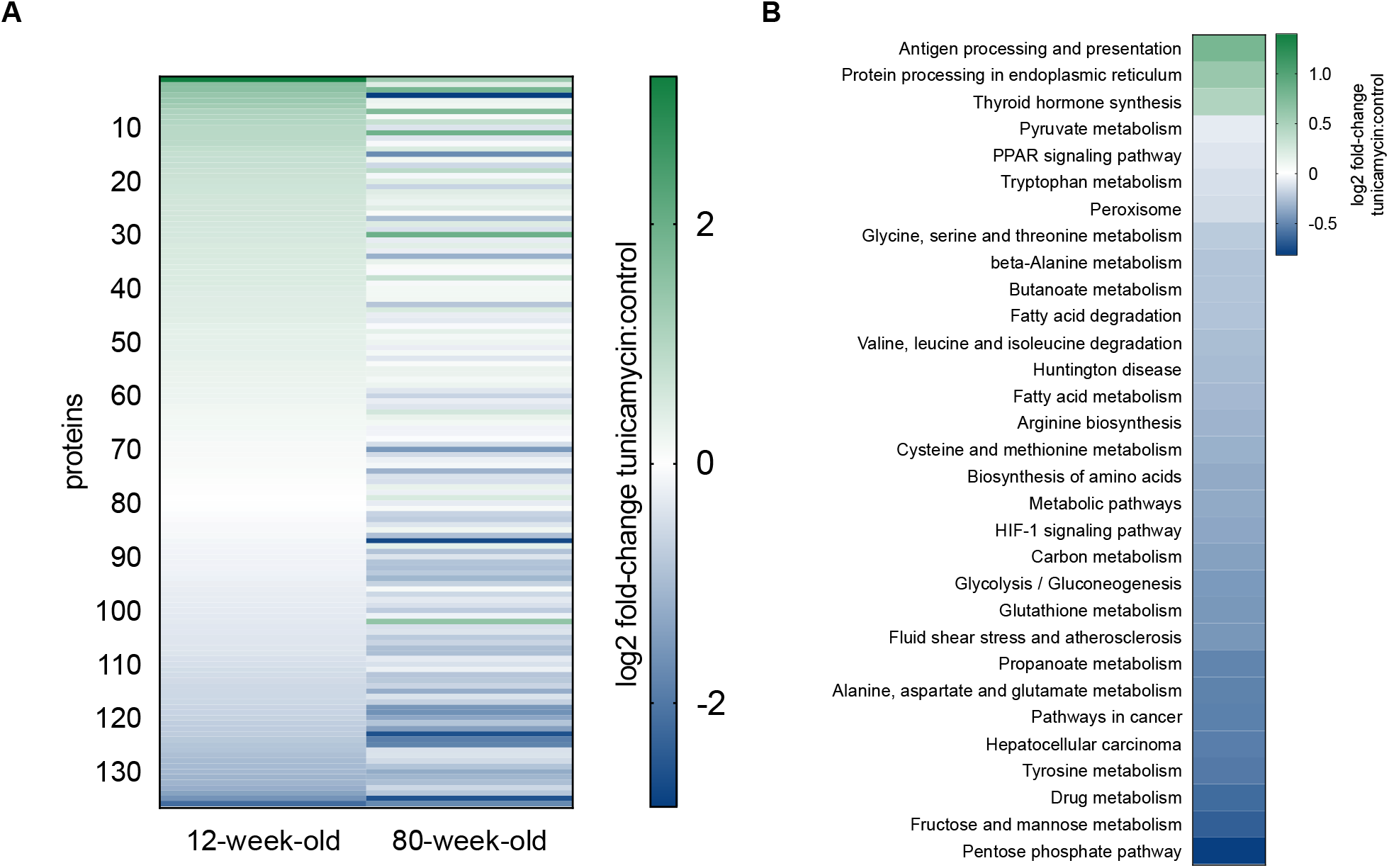
Effects of age on global protein synthesis rates with tunicamycin treatment for paired proteins in young and aged mice. (a) Proteome-wide tunicamycin induced changes in individual protein translation rates (n=136 proteins). Values expressed as log2 fold-change of tunicamycin treated/control. 12-week-old mice values expressed on the left, and protein matched 80-week-old values expressed on the right. (b) KEGG-pathway analysis of the log2 fold-change of 80-week-old-mice treated with tunicamycin to 12-week-old-mice treated with tunicamycin (1=no change). n = at least 5 proteins per pathway.

### BiP synthesis is higher with aging under ER stress conditions

BiP, a key chaperone involved in the UPR, was much more rapidly synthesized in aged mice challenged with tunicamycin as compared to young mice. BiP synthesis increased by ∼2-fold in young mice under ER stress but by more than 3-fold in aged mice under ER stress conditions, showing significant increased translation of BiP in aged tunicamycin challenged compared to young. To compare rates of translation to the amount of mRNA present, *bip* mRNA was measured via RT-qPCR. *bip* mRNA showed no significant difference in young mice challenged with tunicamycin, however, was 9.2-fold higher in aged mice challenged with tunicamycin as compared to control. *xbp1s* mRNA, a spliced version of *xbp1* indicative of initiation of the UPR, was also measured. Young mice showed no significant differences when challenged with tunicamycin, however, aged mice showed a 3.5-fold increase (figure 4). Abundance of BiP was measured via western blot, and no differences between young and aged were seen after challenge with tunicamycin.

**Figure 4:**
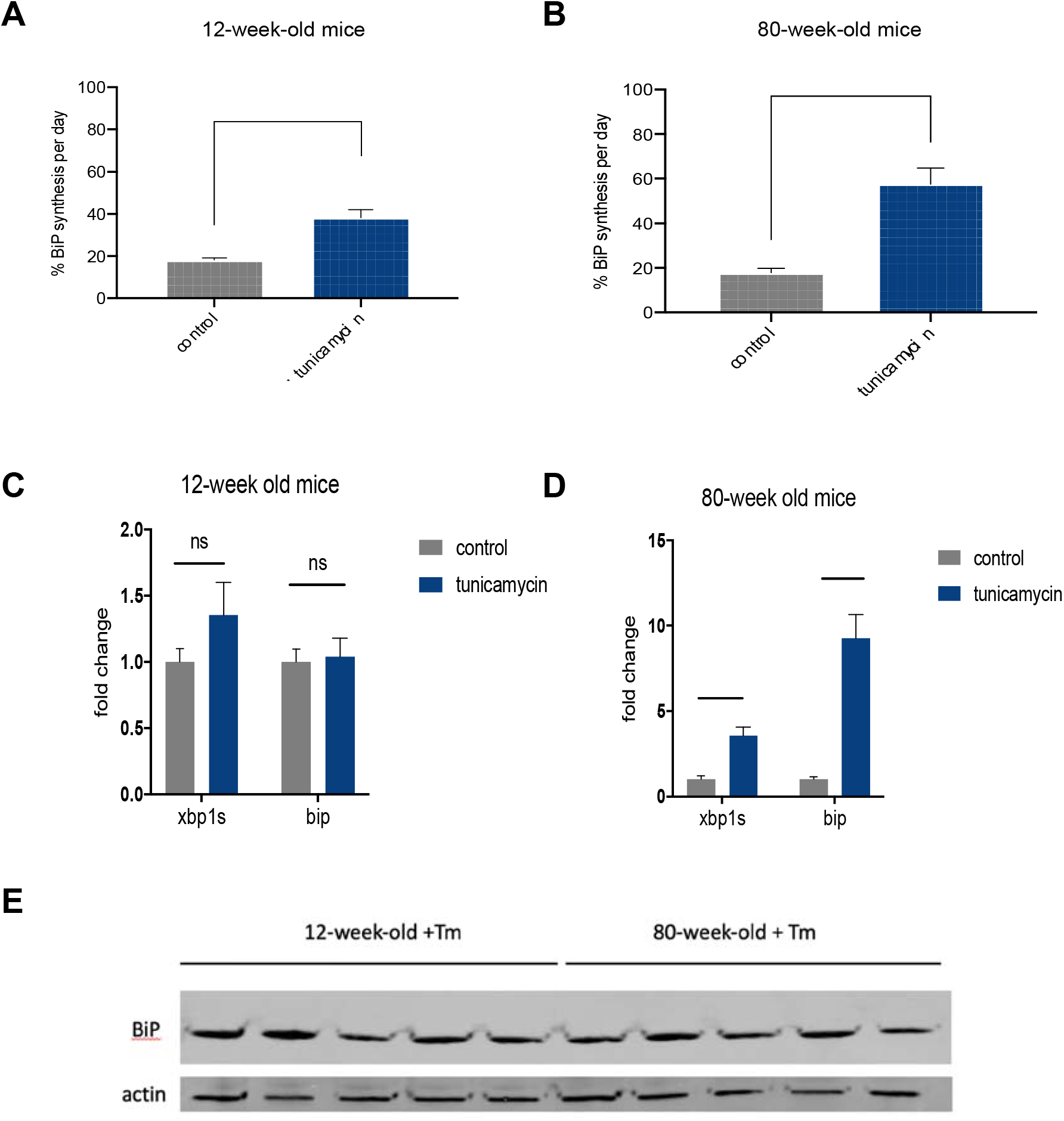
(a) BiP protein synthesis rate per day for 12-week-old mice (n=5). (b) BiP protein synthesis rate per day for 80-week-old mice (n=5). (c) RT-qPCR for *bip* and *xbp1s* in 12-week-old mice (n=5). (d) RT-qPCR for bip and xbp1s in 80-week-old mice (n=5). (e) Western blot for BiP and loading control, actin, for 12-week-old mice vs 80-week-old mice treated with tunicamycin.

### De novo lipogenesis (DNL) is suppressed by both induction of the UPR and age

By quantifying deuterium incorporation into newly synthesized lipids, the contribution from DNL to liver lipids during the treatment period was measured. Rates of DNL were measured for both palmitate incorporated into hepatic phospholipid and triglyceride fractions. Young mice experiencing chronic ER stress showed a significant reduction in *de novo* palmitate incorporation into triglycerides, however, we saw no significant change in DNL contribution to phospholipids. In contrast, aged mice experiencing chronic ER stress displayed a significant reduction in *de novo* palmitate that incorporated into both triglyceride and phospholipid fractions (figure 5).

**Figure 5:**
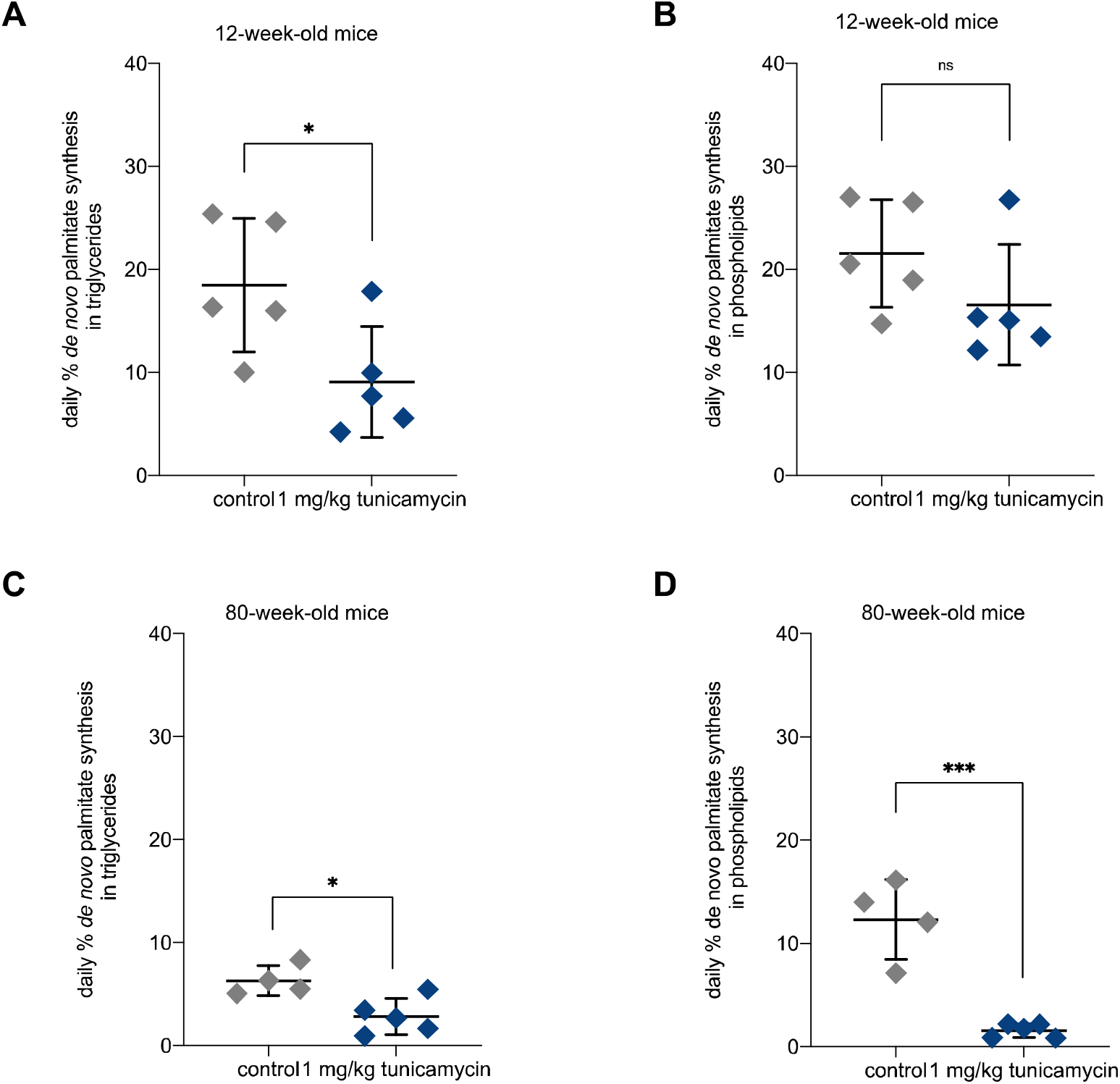
(a) *De novo* lipogenesis (DNL) fractional contribution to palmitate incorporated into triglycerides in 12-week-old control and tunicamycin treated mice (n=5 per group). (b) DNL contribution to palmitate incorporated into phospholipids in 12-week-old control and tunicamycin treated mice (n=5 per group). (c) DNL contribution to palmitate incorporated into triglycerides in 80-week-old control and tunicamycin treated mice (n=5 per group). (d) DNL contribution to palmitate incorporated into phospholipids in 80-week-old control and tunicamycin treated mice (n=5 per group). ns= no significance, * = <0.05, ** = < 0.01, *** = < 0.001.

## Discussion

### Chronic UPR signature in young mice

Young mice challenged with chronic ER stress demonstrated increased translation of proteins involved in *protein processing in the ER*, however, showed modest changes in other ontologies. This suggests that by day 4 of tunicamycin treatment, these mice may be starting to recover from the induced ER stress and are able to handle the load of the tunicamycin. Compared to acute treatment, as previously described^11^, by day 4 translation rates have started to recover however, key UPR^ER^ regulators such as chaperones and BiP remain upregulated. As expected we have previously observed tunicamycin-induced proteome-wide reductions in protein synthesis rates *in vivo* in rodent liver in the hours after single-dose tunicamycin treatment^11^, so the fact that most ontologies are still being translated at normal rates indicates partial recovery. The increase of chaperones and BiP synthesis remained, indicative of an active UPR, however, with suppressed synthesis of proteins involved in lipid metabolism but no reduction in most other ontologies. Interestingly, we found significantly increased BIP synthesis rates despite no change in bip mRNA levels after UPR induction in young mice. This indicates that directly measuring protein synthesis by heavy water labeling with LC/MS-MS analysis may be more sensitive to changes in translational activity than measurement of mRNA levels by RT-qPCR.

### Aging exaggerates UPR metabolic flux signature

Aged mice experiencing chronic ER stress showed a more exaggerated UPR signature as compared to their young counterparts. Aged mice had more suppressed rates of translation across the proteome but sustained the same increase in translation of proteins involved in *protein processing in the ER*, an ontology that encompasses many known UPR proteins such as BiP and protein disulfide isomerases seen in our data set. BiP synthesis was indeed significantly increased in aged mice challenged with tunicamycin as compared to young, which was consistent with *bip* mRNA levels. Abundance of BiP protein by Western blot appeared to be the same when measured in both young and aged mice challenged with tunicamycin, in contrast to higher message levels and synthesis rates increased in aged mice. This may be indicative of more rapid clearance of BiP in aged mice, compensating for higher synthesis rate, or may reflect less sensitivity of the measurement. These data highlight the complex translational control mechanisms used to restore protein homeostasis. Overall, under ER stress we saw lower translation rates of most metabolic protein ontologies, including amino acid metabolism and glucose metabolism. These ontologies were further suppressed in aged mice under ER stress conditions, exemplifying an exaggerated UPR^ER^. A decrease in glutathione metabolism in aged mice under ER stress conditions compared to young mice was seen as well, suggesting a worse dysregulation of redox homeostasis with aging.

### Dysregulation of lipid metabolism

Ontologies involved in lipid metabolism, such as *fatty acid degradation* and *PPAR signaling* were more suppressed in aged mice challenged with tunicamycin, which was consistent with our data showing *de novo* fatty acid synthesis rates were reduced. Aged mice showed a more striking decline in DNL as compared to young mice, specifically in the phospholipid fraction. As phospholipids are the lipid species that comprise membrane, these mice may be worse off in their ability to expand their ER membranes under states of ER stress^24^. Young mice exhibited decline in DNL contribution to palmitate incorporated into triglycerides, but not phospholipids, indicating after 4 days of chronic ER stress they have adapted to be better equipped to handle the accumulation of misfolded proteins in their ER membranes due to continued tunicamycin exposure. Other groups have seen a similar lower DNL phenotype *in vivo* as well^12,13^, indicative of a systemic difference in lipogenic response to UPR^ER^ induction *in vivo* as compared to cellular models^25^ which typically show increased rates of lipogenesis under ER stress. These differences can be reconciled by the ability of a whole organism to draw from other tissues for lipid sources *in vivo*. We have previously shown that lipids are taken up and utilized from other tissues, such as the adipose tissue, under ER stress conditions^11^. Synthesis of phospholipids may be an important part of the UPR due to their ability to be incorporated into the ER membrane and provide raw materials for ER membrane expansion. In yeast, ER membrane expansion has been seen with initiation of the UPR, thus, it might be speculated that this decrease in phospholipid synthesis in aged mice could hinder their ability to recover from ER stress due to spatial constraints^10^.

### Slowed recovery in aged mice challenged with chronic ER stress

We conclude that aging leads to an exaggerated chronic ER stress metabolic signature, including suppressed synthesis of most protein ontologies, with the exception of *protein processing in the ER*, which remains upregulated (figure 6). These data in combination with higher BiP synthesis and dysregulated lipid metabolism in aged mice challenged with tunicamycin indicate that aged mice are not as capable at handling and recovering from ER stress. They sustained high levels of BiP synthesis throughout the 4-day treatment period, whereas young mice showed less increase in BiP synthesis and no significant upregulation of *bip* mRNA. In summary, our data suggests that aging leads to breakdown in the efficiency of the UPR^ER^, leading to an exaggerated UPR metabolic flux signature. This breakdown in the UPR^ER^ with age may be a contributing factor to why diseases manifest as we age. In the liver, specifically, a less capable UPR^ER^ with aging may contribute to metabolic diseases including non-alcoholic fatty liver disease.

**Figure 6:**
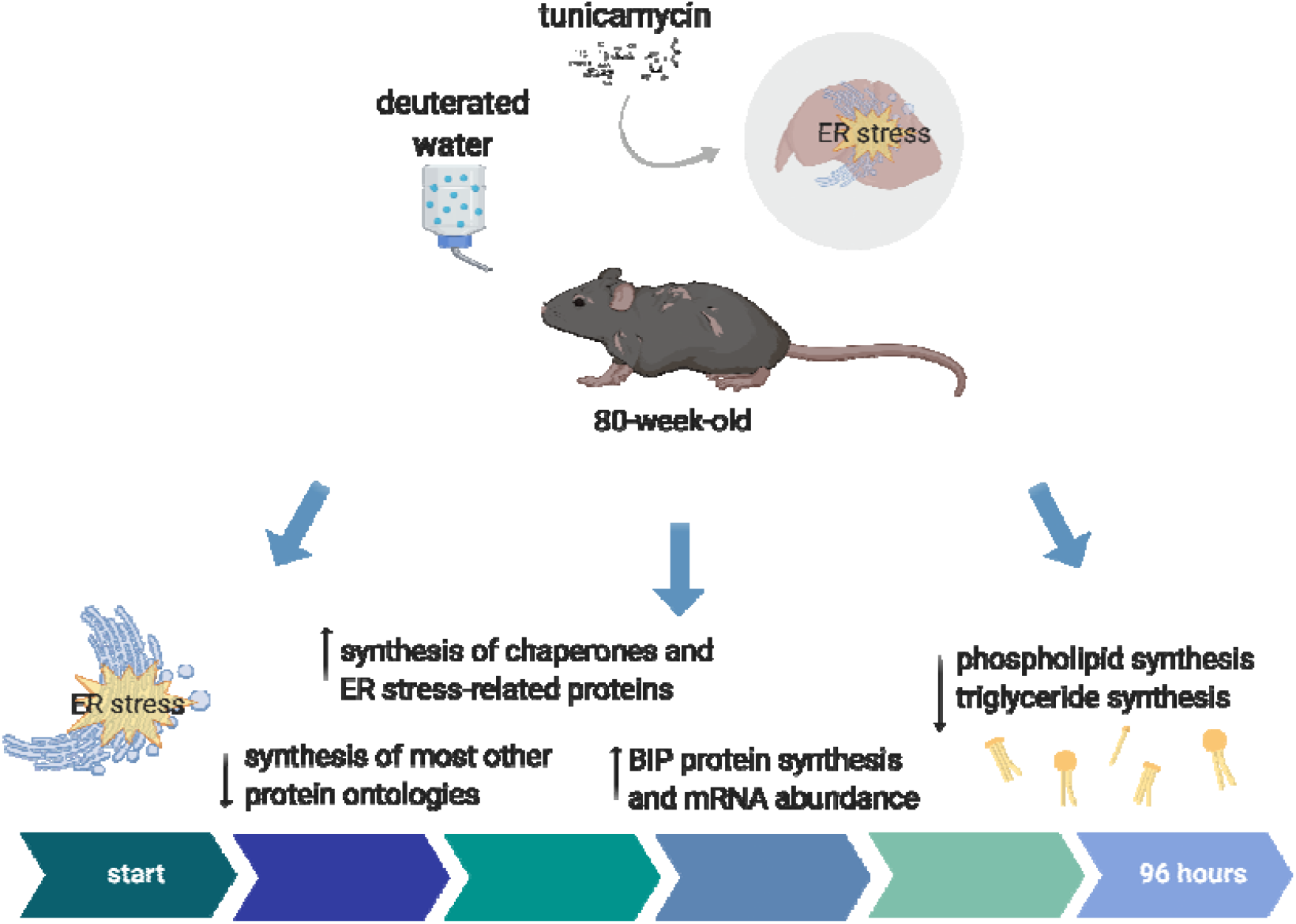
Summary figure. Exaggerated UPR signature in aged animals, with higher translation of ER stress-related proteins, lower translation of all other proteins, lower rates of *de novo* lipogenesis, and BiP protein synthesis and mRNA abundance higher in aged animals with tunicamycin induced ER stress.

## Experimental Procedures

### Animals

C57BL/6J male mice acquired from The Jackson Laboratory were used for this study. Mice were aged to either 12 or 80 weeks. All mice were housed according to the Animal Care and Use Committee (ACUC) standards in the animal facility at UC Berkeley. Mice were fed a standard chow diet and water ad libitum.

### Deuterated water labeling and tunicamycin treatment in mice

Mice were labeled with deuterated water (heavy water, ^2^H_2_O) beginning at time point 0 (t^0^) through the end of the experiment. Proteins synthesized after t^0^ will incorporate deuterium-labeled amino acids, thus enabling the measurement of proteins synthesized during the period of exposure to heavy water. Deuterium is rapidly incorporated throughout the body of an organism after treatment, bringing the deuterium enrichment in body water up to 5%. Deuterium enrichment is maintained through the intake of 8% ^2^H_2_O given as drinking water, thus making it an optimal labeling approach for *in vivo* experimental study. Mice are injected intraperitoneally (IP) with 100% ^2^H_2_O containing either tunicamycin dissolved in DMSO, or DMSO control. Mice were treated at 1 mg/kg tunicamycin one per day, or no drug control, and tissues were harvested 96 hours after injections (n=5 mice per group).

### Body water enrichment analysis

Mouse liver were distilled overnight upside down on a bead bath at 85°C to evaporate out body water. Deuterium present in the body water were exchanged onto acetone, and deuterium enrichment in the body water was measured via gas chromatography mass spectrometry (GC-MS)^26^.

### Tissue preparation for liquid chromatography-mass spectrometry (LC-MS)

Tissues were flash frozen after harvest and homogenized in homogenization buffer (100 mM PMSF, 500 mM EDTA, EDTA-free Protease Inhibitor Cocktail (Roche, catalog number 11836170001), PBS) using a 5 mm stainless steel bead at 30 hertz for 45 seconds in a TissueLyser II (Qiagen). Samples were then centrifuged at 10,000 rcf for 10 minutes at 4°C. The supernatant was saved and protein was quantified using a Pierce BCA protein assay kit (ThermoFisher, catalog number 23225). 100 ug of protein was used per sample. 25 uL of 100 mM ammonium bicarbonate solution, 25 uL TFE, and 2.3 uL of 200 mM DTT were added to each sample and incubated at 60°C for 1 hour. 10 uL 200 mM iodoacetamide was then added to each sample and allowed to incubate at room temperature in the dark for 1 hour. 2 uL of 200 mM DTT was added and samples were incubated for 20 minutes in the dark. Each sample was then diluted with 300 uL H_2_O and 100 uL 100 mM ammonium bicarbonate solution. Trypsin was added at a ratio of 1:50 trypsin to protein (trypsin from porcine pancreas, Sigma Aldrich, catalog number T6567). Samples were incubated at 37°C overnight. The next day, 2 uL of formic acid was added. Samples were centrifuged at 10,000 rcf for 10 minutes, collecting the supernatant. Supernatant was speedvac’d until dry and re-suspended in 50 uL of 0.1 % formic acid/3% acetonitrile/96.9% LC-MS grade water and transferred to LC-MS vials to be analyzed via LC-MS.

### Liquid chromatography-mass spectrometry (LC-MS) analysis

Trypsin-digested peptides were analyzed on a 6550 quadropole time of flight (Q-ToF) mass spectrometer equipped with Chip Cube nano ESI source (Agilent Technologies). High performance liquid chromatography (HPLC) separated the peptides using capillary and nano binary flow. Mobile phases were 95% acetonitrile/0.1% formic acid in LC-MS grade water. Peptides were eluted at 350 nl/minute flow rate with an 18 minute LC gradient. Each sample was analyzed once for protein/peptide identification in data-dependent MS/MS mode and once for peptide isotope analysis in MS mode. Acquired MS/MS spectra were extracted and searched using Spectrum Mill Proteomics Workbench software (Agilent Technologies) and a mouse protein database (www.uniprot.org). Search results were validated with a global false discovery rate of 1%. A filtered list of peptides was collapsed into a nonredundant peptide formula database containing peptide elemental composition, mass, and retention time. This was used to extract mass isotope abundances (M0-M3) of each peptide from MS-only acquisition files with Mass Hunter Qualitative Analysis software (Agilent Technologies). Mass isotopomer distribution analysis (MIDA) was used to calculate peptide elemental composition and curve-fit parameters for predicting peptide isotope enrichment based on precursor body water enrichment (p) and the number (n) of amino acid C-H positions per peptide actively incorporating hydrogen (H) and deuterium (D) from body water. Subsequent data handling was performed using python-based scripts, with input of precursor body water enrichment for each subject, to yield fractional synthesis rate (FSR) data at the protein level. FSR data were filtered to exclude protein measurements with fewer than 2 peptide isotope measurements per protein. Details of FSR calculations and data filtering criteria were described previously^19^.

### Calculation of fractional replacement (f) and replacement rate constant (k) for individual proteins

Details of f calculations were previously described^15^. These values were used to generate the ratio of tunicamycin treated to untreated synthesis rates.

### Statistical analysis

Data were analyzed using GraphPad Prism software (version 8.0).

### Tissue preparation for gas chromatography-mass spectrometry (GC-MS)

A chloroform methanol extraction was used to isolate lipids from the liver tissue. These lipids were run on a thin-layer chromatography (TLC) plate to separate phospholipid and triglyceride fractions. These fractions containing the palmitate were further derivatized for GC-MS analysis

### Gas chromatography-mass spectrometry (GC-MS) analysis

Palmitate isotopic enrichments were measured by GC-MS (Agilent models 6890 and 5973; Agilent, Inc., Santa Clara, CA) using helium carrier gas, a DB-225 (DB 17 for cholesterol and DB 225 for palmitates) fused silica column (30M x 0.25mm ID x 0.25um), electron ionization mode, and monitoring m/z 385, 386, and 387 for palmitates for M0, M1, and M2 respectively, as previously described^21^. Palmitate methyl ester enrichments were determined by GC-MS using a DB-17 column (30M x 0.25mm ID x 0.25um), with helium as carrier gas, electron ionization mode, and monitoring m/z 270, 271, and 272 for M0, M1, and M2. Baseline unenriched standards for both analytes were measured concurrently to correct for abundance sensitivity.

### Calculation of de novo lipogenesis (DNL)

The measurement of newly synthesized fatty acids formed during ^2^H_2_O labeling period was assessed using a combinatorial model of polymerization biosynthesis, as described previously^21^. Mass isotopomer distribution analysis (MIDA) along with body ^2^H_2_O enrichment, representing the precursor pool enrichment (p), is used to determine the theoretical maximum enrichment of each analyte. Using the measured deuterium enrichments, fractional and absolute contributions from DNL are then calculated. The value for f DNL represents the fraction of total triglyceride or phospholipid palmitate in the depot derived from DNL during the labeling period, and absolute DNL represents grams of palmitate synthesized by the DNL pathway.

### Western blot

Starting with frozen tissue, tissue was homogenized in homogenization buffer (100 mM PMSF, 500 mM EDTA, EDTA-free Protease Inhibitor Cocktail (Roche, catalog number 11836170001), PBS) using a 5 mm stainless steel bead at 30 hertz for 45 seconds in a TissueLyser II (Qiagen). Samples were then centrifuged at 10,000 rcf for 10 minutes at 4C. The supernatant was saved and protein was quantified using a Pierce BCA protein assay kit (ThermoFisher, catalog number 23225). 30 ug of protein was used per sample. 2X Laemmli sample buffer was added (Sigma, catalog number S3401) at a 1:1 ratio. Samples were brought to the same volume with 1% SDS, vortexed briefly, and heated in a heating block for 10 minutes at 95°C. Samples were tip sonicated for 10 seconds and then centrifuged for 5 minutes at 15,000 g. 4-12% gradient polyacrylamide gels were used with MES buffer and gels were run at 120V until loading dye line passed through gel. iBlot2 was used to transfer the gel onto a PVDF membrane. Membranes were washed 3 times with PBST and then blocked with 5% milk for 1 hour. Membranes were then washed again with PBST 3 times. BiP (Cell Signaling Technology, catalog number 3183S) and actin (Santa Cruz technology, catalog number 47778) antibodies were diluted in 5% BSA and rotated at 4°C overnight. Membranes were then washed 3 times with PBST, and LiCor secondary antibodies diluted in 5% BSA were added and rotated for 2 hours at room temperature. Membranes were washed 3 times with PBST and then imaged using a LiCor imaging system.

### Quantitative reverse transcription PCR (RT-qPCR)

RNA was isolated using standard Trizol protocol and RNA concentrations were obtained using a Nanodrop. After normalizing concentrations, cDNA was synthesized using 2 ug RNA with RevertAid RT Kit (Thermofisher, catalog number K1691). Maxima SYBR Green/ROX qPCR Master Mix (ThermoFisher, catalog number K0221) was used for RT-qPCR. Actin was used to normalize. Oligonucleotide sequences used:

> bip F: CGAGGAGGAGGACAAGAAGG
>
> bip R: CACCTTGAACGGCAAGAACT
>
> xbp1s forward: TGCTGAGTCCGCAGCAGGTG
>
> xbp1s reverse: GCTGGCAGGCTCTGGGGAAG
>
> actin forward: CGAATCATGAGCATTGTAGAC
>
> actin reverse: GTAATTCTTATCTCCAGCCAG

### KEGG pathway analysis

Protein fractional synthesis rates were weighted by the peptide count and averaged according to their KEGG pathway involvements. We used the Uniprot.ws package in R from Bioconductor to find mappings between UniProt accession numbers and their corresponding KEGG IDs for each protein. Tables were generated for the entire known proteome for mouse. We then used the Bio.KEGG module of Biopython in Python to access to the REST API of the KEGG database to get a list of pathways to which each protein belongs. A set of all the pathways relevant to the experiment was generated and each protein and its corresponding fold change value were assigned to each pathway. KEGG pathways with no less than five proteins were used for representation of the data.

## Acknowledgments

The authors would like to thank Marcy Matthews, Mark Fitch, and the entire Hellerstein lab for their technical assistance and feedback. We would also like to thank the Dillin lab at UC Berkeley for insightful input. Thanks to Monica Forsythe and all of the support staff at the UC Berkeley Animal Facility for assistance with mouse injections and animal care and housing.

Graphical abstract and figures generated with BioRender. We thank Fred Ward for miscellaneous support.

## Author contributions

CPS, AD, and MKH conceived and designed experiments. CPS, LP, SY, and JH performed experiments. HP assisted with tunicamycin treatment in mice. HM and EN assisted with LC-MS and GC-MS sample processing and data analysis. NZ performed KEGG pathway analysis. CPS, AEF, and MKH contributed to writing the manuscript.

## Figures

**Supplementary Figure 1.**
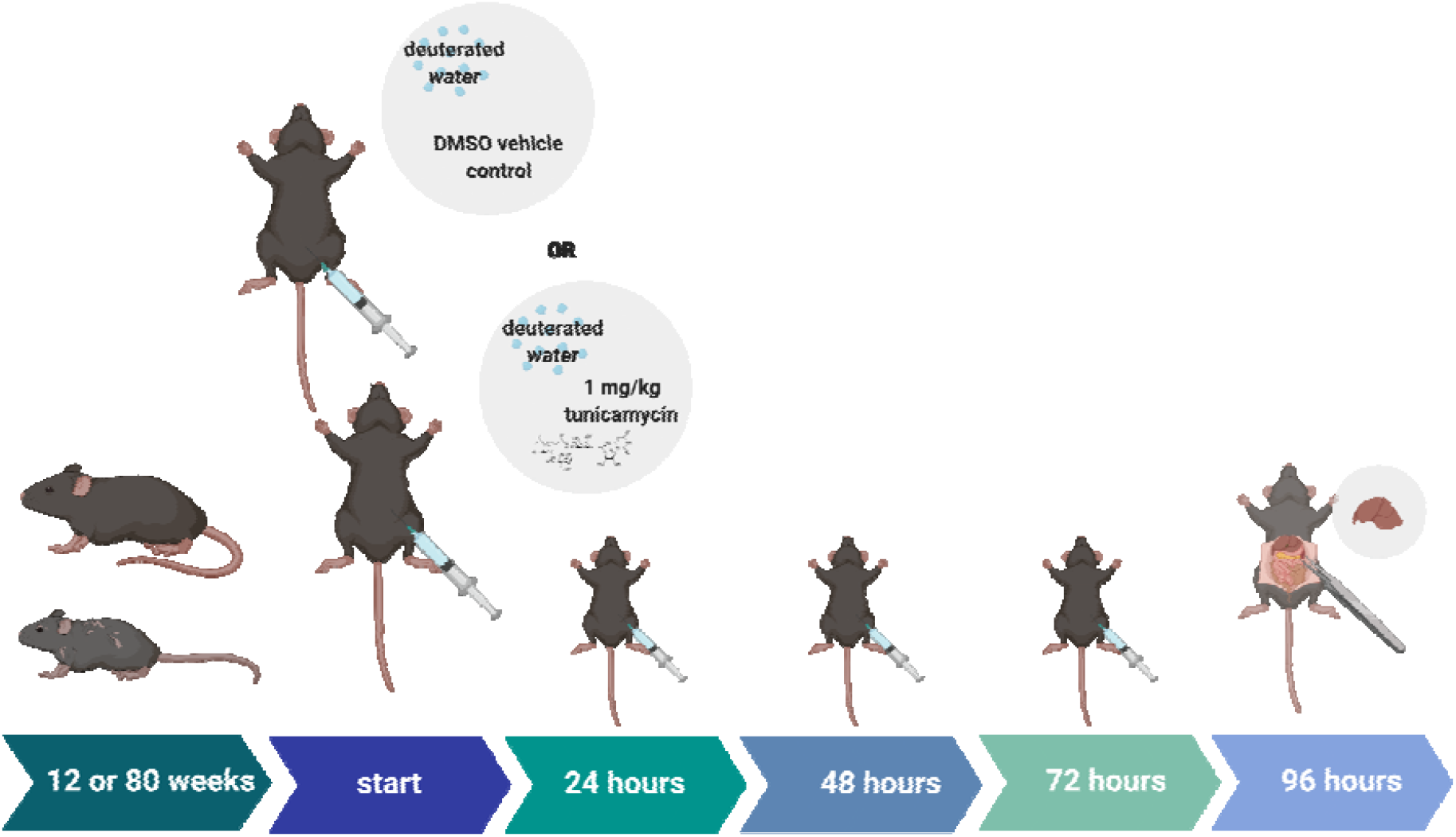
Study design: 12-week-old or 80-week-old mice (n=5 per group) were treated with 1 mg/kg tunicamycin or vehicle control, DMSO, once per day for 3 days. Mice were injected with 35uL/g deuterated water on day 1. Mice were sacrificed and livers were taken on day 4.

**Supplementary Table 1:** Individual protein synthesis rate ratios of aged control mice treated with tunicamycin. Ratio of above 1 indicates a higher synthesis rate with age and/or tunicamycin treatment. Ratio of below 1 indicates a lower synthesis rate with age and/or tunicamycin treatment.

| **Protein** | **ratio aged**<br>**tm/control** | **ratio young**<br>**tm/control** | **ratio aged**<br>**tm/young tm** |
| --- | --- | --- | --- |
| 10 kDa heat shock protein, mitochondrial | 1.21 | 1.11 | 1.10 |
| 2-iminobutanoate/2-iminopropanoate deaminase | 0.83 | 1.00 | 0.83 |
| 3-hydroxyacyl-CoA dehydrogenase type-2 | 0.35 | 1.06 | 0.33 |
| 3-ketoacyl-CoA thiolase A, peroxisomal | 1.41 | 1.00 | 1.41 |
| 3-ketoacyl-CoA thiolase B, peroxisomal | 0.76 | 0.79 | 0.95 |
| 3-ketoacyl-CoA thiolase, mitochondrial | 0.76 | 0.89 | 0.85 |
| 40S ribosomal protein S4, X isoform | 1.42 | 1.31 | 1.08 |
| 60 kDa heat shock protein, mitochondrial | 1.10 | 1.19 | 0.92 |
| 60S ribosomal protein L29 | 0.90 | 1.08 | 0.83 |
| 60S ribosomal protein L6 | 1.72 | 1.38 | 1.24 |
| Actin, alpha skeletal muscle | 1.17 | 1.19 | 0.98 |
| Actin, cytoplasmic 2 | 1.11 | 1.11 | 1.00 |
| Acyl-CoA synthetase family member 2, mitochondrial | 0.43 | 0.48 | 0.88 |
| Acyl-CoA-binding protein | 0.57 | 0.69 | 0.83 |
| Acyl-coenzyme A synthetase ACSM1, mitochondrial | 1.05 | 1.50 | 0.70 |
| Adenosylhomocysteinase | 1.10 | 1.37 | 0.80 |
| Alanine--glyoxylate aminotransferase 2, mitochondrial | 0.30 | 1.69 | 0.18 |
| Alcohol dehydrogenase 1 | 0.43 | 0.67 | 0.64 |
| Alcohol dehydrogenase class-3 | 0.58 | 0.78 | 0.74 |
| Aldehyde dehydrogenase family 8 member A1 | 0.96 | 1.39 | 0.69 |
| Aldehyde dehydrogenase, cytosolic 1 | 0.64 | 1.17 | 0.55 |
| Aldehyde dehydrogenase, mitochondrial | 0.73 | 0.79 | 0.92 |
| Alpha-1-antitrypsin 1-3 | 0.84 | 1.41 | 0.60 |
| Alpha-enolase | 0.77 | 0.94 | 0.81 |
| Arginase-1 | 0.77 | 1.13 | 0.68 |
| Argininosuccinate synthase | 1.11 | 1.34 | 0.83 |
| Aspartate aminotransferase, mitochondrial | 0.80 | 1.30 | 0.62 |
| ATP synthase subunit alpha, mitochondrial | 0.73 | 1.02 | 0.72 |
| ATP synthase subunit beta, mitochondrial | 0.69 | 1.06 | 0.66 |
| Betaine--homocysteine S-methyltransferase 1 | 0.69 | 0.50 | 1.38 |
| Bifunctional epoxide hydrolase 2 | 0.78 | 1.24 | 0.63 |
| Calreticulin | 3.79 | 1.88 | 2.01 |
| Carbamoyl-phosphate synthase [ammonia], mitochondrial | 0.64 | 0.97 | 0.66 |
| Carbonic anhydrase 2 | 0.15 | 0.91 | 0.17 |
| Carbonic anhydrase 3 | 0.17 | 0.33 | 0.51 |
| Carboxylesterase 1D | 0.26 | 0.57 | 0.46 |
| Carboxylesterase 1F | 0.34 | 0.65 | 0.52 |
| Carboxylesterase 3A | 0.38 | 0.60 | 0.63 |
| Carboxylesterase 3B | 0.56 | 0.39 | 1.45 |
| Catalase | 0.74 | 0.72 | 1.03 |
| Clathrin heavy chain 1 | 0.86 | 1.04 | 0.82 |
| Cystathionine gamma-lyase | 1.13 | 2.34 | 0.48 |
| Cysteine sulfinic acid decarboxylase | 0.16 | 0.59 | 0.28 |
| Cytochrome c oxidase subunit 6A1, mitochondrial | 1.22 | 1.00 | 1.21 |
| Cytochrome P450 2D10 | 0.94 | 1.85 | 0.51 |
| Cytosolic 10-formyltetrahydrofolate dehydrogenase | 0.53 | 0.93 | 0.57 |
| D-dopachrome decarboxylase | 1.19 | 1.20 | 0.99 |
| Delta-1-pyrroline-5-carboxylate dehydrogenase, mitochondrial | 0.65 | 0.68 | 0.96 |
| Electron transfer flavoprotein subunit alpha, mitochondrial | 0.97 | 1.08 | 0.90 |
| Electron transfer flavoprotein subunit beta | 0.84 | 1.16 | 0.73 |
| Elongation factor 1-alpha 1 | 1.29 | 1.41 | 0.91 |
| Elongation factor 2 | 0.86 | 0.70 | 1.22 |
| Endoplasmic reticulum chaperone BiP | 3.22 | 2.11 | 1.53 |
| Endoplasmin | 3.53 | 2.92 | 1.21 |
| Estradiol 17 beta-dehydrogenase 5 | 0.32 | 0.64 | 0.50 |
| Fatty acid synthase | 0.93 | 1.61 | 0.58 |
| Fatty acid-binding protein, liver | 0.52 | 0.47 | 1.12 |
| Ferritin light chain 1 | 1.10 | 1.25 | 0.88 |
| Formimidoyltransferase-cyclodeaminase | 1.39 | 1.75 | 0.79 |
| Fructose-1,6-bisphosphatase 1 | 1.33 | 1.50 | 0.88 |
| Fructose-bisphosphate aldolase B | 0.84 | 1.31 | 0.64 |
| Fumarylacetoacetase | 0.75 | 1.02 | 0.74 |
| Glutamate dehydrogenase 1, mitochondrial | 0.55 | 0.61 | 0.89 |
| Glutathione peroxidase 1 | 0.65 | 0.78 | 0.83 |
| Glutathione S-transferase A1 | 0.52 | 0.76 | 0.69 |
| Glutathione S-transferase A3 | 0.47 | 0.87 | 0.54 |
| Glutathione S-transferase Mu 1 | 1.17 | 2.50 | 0.47 |
| Glutathione S-transferase Mu 3 | 1.34 | 2.95 | 0.46 |
| Glutathione S-transferase P 1 | 0.52 | 0.77 | 0.68 |
| Glyceraldehyde-3-phosphate dehydrogenase | 0.85 | 1.26 | 0.67 |
| Glycine N-methyltransferase | 1.26 | 1.53 | 0.83 |
| Glycogen phosphorylase, liver form | 0.56 | 0.70 | 0.80 |
| Glyoxylate reductase/hydroxypyruvate reductase | 1.11 | 1.33 | 0.83 |
| Heat shock cognate 71 kDa protein | 2.75 | 0.80 | 3.44 |
| Heat shock protein HSP 90-beta | 0.75 | 1.47 | 0.51 |
| Hemoglobin subunit alpha | 0.60 | 0.81 | 0.73 |
| Hemoglobin subunit beta-1 | 0.63 | 0.85 | 0.74 |
| Homogentisate 1,2-dioxygenase | 0.50 | 1.50 | 0.34 |
| Hydroxyacyl-coenzyme A dehydrogenase, mitochondrial | 0.47 | 0.47 | 1.01 |
| Hydroxymethylglutaryl-CoA lyase, mitochondrial | 1.07 | 0.99 | 1.08 |
| Hydroxymethylglutaryl-CoA synthase, mitochondrial | 0.96 | 1.28 | 0.75 |
| Isochorismatase domain-containing protein 2A | 0.77 | 1.88 | 0.41 |
| Isocitrate dehydrogenase [NADP] cytoplasmic | 1.07 | 0.84 | 1.27 |
| L-lactate dehydrogenase A chain | 0.86 | 1.00 | 0.86 |
| Major urinary protein 1 | 0.55 | 0.89 | 0.61 |
| Major urinary protein 17 | 0.67 | 0.84 | 0.80 |
| Major urinary protein 2 | 0.46 | 1.03 | 0.44 |
| Malate dehydrogenase, cytoplasmic | 1.05 | 1.39 | 0.76 |
| Malate dehydrogenase, mitochondrial | 1.32 | 1.59 | 0.83 |
| Maleylacetoacetate isomerase | 1.20 | 1.21 | 0.99 |
| Medium-chain specific acyl-CoA dehydrogenase, mitochondrial | 0.44 | 1.41 | 0.31 |
| Methanethiol oxidase | 0.75 | 0.55 | 1.37 |
| Methylmalonate-semialdehyde dehydrogenase [acylating], mitochondrial | 1.16 | 1.27 | 0.92 |
| Microsomal glutathione S-transferase 1 | 0.80 | 0.84 | 0.96 |
| NADP-dependent malic enzyme | 0.75 | 0.52 | 1.44 |
| Non-specific lipid-transfer protein | 0.58 | 0.88 | 0.65 |
| Nucleoside diphosphate kinase B | 1.48 | 1.11 | 1.33 |
| Ornithine carbamoyltransferase, mitochondrial | 0.75 | 0.67 | 1.11 |
| Peptidyl-prolyl cis-trans isomerase A | 0.89 | 1.10 | 0.81 |
| Peroxiredoxin-1 | 0.78 | 1.17 | 0.67 |
| Peroxiredoxin-6 | 0.81 | 0.74 | 1.10 |
| Peroxisomal acyl-coenzyme A oxidase 1 | 1.34 | 1.54 | 0.87 |
| Phenylalanine-4-hydroxylase | 1.27 | 1.27 | 0.99 |
| Phosphoglucomutase-1 | 0.76 | 1.89 | 0.40 |
| Phosphoglycerate kinase 1 | 1.21 | 1.40 | 0.87 |
| Phosphoglycerate mutase 1 | 0.73 | 0.82 | 0.90 |
| Polycystic kidney disease protein 1-like 1 | 0.30 | 0.22 | 1.35 |
| Pregnancy zone protein | 1.18 | 1.52 | 0.77 |
| Protein disulfide-isomerase | 1.66 | 1.99 | 0.83 |
| Protein disulfide-isomerase A3 | 1.86 | 1.62 | 1.15 |
| Protein disulfide-isomerase A4 | 3.90 | 1.46 | 2.67 |
| Protein disulfide-isomerase A6 | 2.51 | 9.38 | 0.27 |
| Protein/nucleic acid deglycase DJ-1 | 0.94 | 1.38 | 0.68 |
| Pyruvate carboxylase, mitochondrial | 0.62 | 0.66 | 0.94 |
| Regucalcin | 0.28 | 0.55 | 0.51 |
| Retinal dehydrogenase 1 | 0.59 | 0.97 | 0.61 |
| S-adenosylmethionine synthase isoform type-1 | 1.36 | 1.48 | 0.92 |
| S-formylglutathione hydrolase | 1.06 | 2.05 | 0.52 |
| S-methylmethionine--homocysteine S-methyltransferase BHMT2 | 0.54 | 0.50 | 1.09 |
| Sarcosine dehydrogenase, mitochondrial | 1.24 | 0.90 | 1.38 |
| SEC14-like protein 2 | 0.88 | 0.80 | 1.09 |
| Selenium-binding protein 2 | 0.67 | 0.43 | 1.54 |
| Serum albumin | 0.40 | 0.63 | 0.64 |
| Short-chain specific acyl-CoA dehydrogenase, mitochondrial | 0.53 | 0.89 | 0.60 |
| Sorbitol dehydrogenase | 0.56 | 1.32 | 0.42 |
| Succinate--CoA ligase [ADP/GDP-forming] subunit alpha, mitochondrial | 0.14 | 2.61 | 0.05 |
| Superoxide dismutase [Cu-Zn] | 0.72 | 1.07 | 0.67 |
| Transketolase | 0.68 | 1.68 | 0.40 |
| Trifunctional enzyme subunit alpha, mitochondrial | 0.92 | 1.04 | 0.89 |
| Triokinase/FMN cyclase | 0.54 | 0.90 | 0.60 |
| Triosephosphate isomerase | 0.84 | 1.45 | 0.57 |
| Tubulin beta-2B chain | 1.09 | 1.27 | 0.86 |
| Tubulin beta-4B chain | 1.04 | 1.21 | 0.86 |
| Valacyclovir hydrolase | 1.13 | 0.94 | 1.20 |
| Xanthine dehydrogenase/oxidase | 1.12 | 1.68 | 0.67 |
| Xylulose kinase | 0.65 | 1.56 | 0.41 |

## Notes

### Competing Interest Statement

The authors have declared no competing interest.

